# The percentage of Monocytes CD39+ is higher in Pregnant COVID-19 than in Non-Pregnant COVID-19 patients

**DOI:** 10.1101/2021.06.18.449054

**Authors:** A. Cérbulo-Vázquez, M. García-Espinosa, J.C. Briones-Garduño, L. Arriaga-Pizano, E. Ferat-Osorio, B. Zavala-Barrios, G.L. Cabrera-Rivera, P. Miranda-Cruz, M.T. García de la Rosa, J.L. Prieto-Chávez, V. Rivero-Arredondo, R.L. Madera-Sandoval, A. Cruz-Cruz, E. Salazar-Rios, ME Salazar-Rios, D Serrano-Molina, R. C. De Lira-Barraza, A. H. Villanueva-Compean, A. Esquivel-Pineda, R. Ramirez-Montes de Oca, F. Caldiño-Soto, L.A. Ramírez-García, G. Flores-Padilla, O. Moreno-Álvarez, GML Guerrero-Avendaño, C. López-Macías

## Abstract

Current medical guidelines consider COVID-19 pregnant women a high-risk group. Physiological gestation down regulates the immunological response to maintain “maternal-fetal tolerance”; hence, a SARS-CoV-2 infection constitutes a potentially threatening condition to both the mother and the fetus. To establish the immune profile in pregnant COVID-19+ patients a cross-sectional study was conducted. Leukocyte immunophenotype, mononuclear leukocyte response to polyclonal stimulus and cytokine/chemokine serum concentration were analyzed in pregnant fifteen COVID-19+ and control groups (fifteen non-pregnant COVID-19+, and thirteen pregnant COVID-19-women). Pregnant COVID-19+ patients exhibit lower percentages of monocytes HLA-DR+ compared with control groups. Nevertheless, pregnant COVID-19+ women show a higher percentage of monocytes CD39+ than controls. Furthermore, a higher concentration of TNF-α, IL-6, MIP1b and IL-4 was observed within the pregnant COVID-19+ group. Our result shows that pregnant women express immunological characteristics that potentially mediate the immune response in COVID-19.

## Introduction

The Severe Acute Respiratory Syndrome Coronavirus (SARS-CoV) and Middle East Respiratory Syndrome Coronavirus (MERS-CoV) infections could result in a high mortality among pregnant women (25% and 27%, respectively) (1). In 2019, a new coronavirus called SARS-CoV-2 appeared. Viral infections lead to a powerful cell and humoral immune response in gestating women, increasing the embryo/fetal-mother morbidity and mortality (2-4). The immune response in pregnant women is mediated by a diverse number of cellular and humoral mechanisms (5, 6) that result in a unique biological scenario that these women face when infected with SARS-CoV-2. Moreover, comorbidities highly prevalent in Mexico, such as obesity, hypertension and diabetes are associated with critical disease in both general population and pregnant women (3, 4, 7, 8). However, the clinical presentation and symptoms are quite similar between the general population and pregnant women COVID-19+ (3, 9).

In contrast to the epidemiological and clinical evaluation, the immune profile of pregnant women with COVID-19 has been poorly explored. Similar to the general population, deep lymphopenia is reported in pregnant women with COVID-19 (10). Other parameters, such as neutrophil count, are useful for a COVID-19 diagnosis (11). Furthermore, an increased number of neutrophil/lymphocyte ratio has been associated to a fatal outcome in COVID-19 patients (12-15). Several studies indicate that leukocyte count is necessary for the initial evaluation of COVID-19 patients. However, a deeper analysis of these leukocytes could improve our knowledge of the SARS-CoV2 infection, especially in pregnant women.

Cytokines and chemokines are key regulators in the coronavirus infection. SARS-CoV induces low expression of IFN-α, IFN-β and IL-10, as well as a moderate expression of TNF-α and IL-6 and a high expression of CCL3, CCL5, CCL-2 and CXCL10 (16) (17). In addition, SARS-CoV’s protein S induces CCL2 and CXCL8 synthesis *in vitro* (18, 19). Also, several humoral components of the immune response are involved in COVID-19, among them, inflammatory cytokine/chemokines which have been observed in high serum concentrations at the general population (20, 21). COVID-19 patients show a higher concentration of IL-2, IL-7, IL-10, G-CSF, IP10 (CXCL-10), MCP-1 (CCL2), MIP1a (CCL3) and TNF-α compared to non COVID-19 patients (20). The source of these cytokine/chemokines is diverse, and could include lung epithelial cells, endothelium cells, and leukocytes (22-24).

Exacerbated inflammatory response in pregnancy could disturb the delicate immune balance, leading to a significant increased morbidity and mortality. To analyze the immune profile in pregnant women with COVID-19+, a cross sectional study was executed. Our analysis includes a) immunophenotype of lymphocytes and monocytes expressing certain activation markers, b) serum cytokine/chemokine concentration, and total blood challenged against polyclonal stimulus, in presence or absence of Brefeldin-A, after 4 hours of culture, c) inflammatory cytokines determination, and d) supernatant cytokine/chemokine concentration. A more detailed picture of the immune map in pregnant women with COVID-19 could help better understand the pathophysiology of SARS-CoV-2 infection and improve the quality of healthcare provided to these women and newborns.

## Material and methods

### Patients

This study was conducted by the “Servicio de Ginecología y Obstetricia” at the Hospital General de Mexico, “Dr. Eduardo Liceaga” in conjunction with the Unidad de Investigación Médica en Inmunoquímica (UIMIQ), at the Hospital de Especialidades, Centro Médico Nacional Siglo XXI, and Servicio de complicaciones de la segunda mitad del embarazo, División Obstetricia. UMAE Hospital de GinecoObstetricia No. 4 “Dr. Luis Castelazo Ayala” (Research project: *DI/20112/04/45, and R-2020-785-095* respectively). After obtaining a signed informed consent letter, forty-four women were enrolled. Three groups were analyzed: a) Non-pregnant COVID-19 positive (NP-COVID-19+, n=15), b) Pregnant COVID-19 positive (P-COVID-19+, n=16), and c) Pregnant COVID-19 negative (P-COVID-19-, n=13). SARS-CoV2 viral infection was confirmed by specific reverse transcription–polymerase chain reaction (RT-PCR). The COVID-19 diagnosis was based on clinical characteristics (25); comorbidities and clinical signs or symptoms were registered and shown in Table 1 and 2 respectively.

**Table 1.** Clinical and laboratory characteristics.

|  | **NP-COVID-19 +**<br>**(n=9-15)a** | **P-COVID-19**<br>**+**<br>**(n=8-15)b** | **P-COVID-19**<br>**-**<br>**(n=13)c** | ***p*** |
| --- | --- | --- | --- | --- |
| **Age (years)** | 34.1±7 | 28±7.1 | 25.6±5.7 | a vs. c<br>0.006 |
| **BMI** | 31.6±6.9 | 30.0±6.3 | 28.0±3.5 | 0.356 |
| **Respiratory Rate**<br>**(breaths per min)** | 23±3.8 | 22.1±6.8 | 18.6±1.1 | a vs. c<br>0.005 |
| **Heart Rate**<br>**(beats per min)** | 91.2±25.1 | 97.3±19.6 | 77.7±7.5 | **b vs. c**<br>**0.048** |
| **Temperature (°C)** | 36.9±1.2 | 36.7±0.7 | 36.3±0.2 | 0.081 |
| **Mean Arterial**<br>**Pressure (mmHg)** | 159.5±17.4 | 153.2±15.4 | 157.1±13.3 | 0.631 |
| **Hemoglobin** | 13±2.7 | 12.6±2.0 | 11.9±1.1 | 0.070 |
| **Leukocytes**<br>**(10xmm<sup>3</sup>)** | 8.6±4 | 11.1±3.2 | 8.4±2.3 | 0.140 |
| **NLR** | 11.8±11.7 | 5.1±1.9 | 3.7±1.2 | a vs. c<br>0.007 |
| **PLT (10xmm<sup>3</sup>)** | 272.3±99.2 | 327.2±218.1 | 232.8±45.4 | 0.548 |
| **Glucose (mg/dL)** | 122.5±51.8 | 91.1±20.1 | 89.3±12.5 | 0.170 |
| **Urea (mg/dL)** | 30.9±20.4 | 17.4±5.7 | 18.3±5.9 | **a vs. b**<br>**0.040**<br>a vs. c<br>0.027 |
| **Creatinin (mg/dL)** | 0.7±0.3 | 1.0±1.1 | 0.5±0.2 | a vs. c<br>0.009 |
| **LDH (U/L)** | 804.9±1389 | 582.8±510.8 | 269.7±90.4 | **b vs. c**<br>**0.005** |
| **PT (seconds)** | 15.2±3.3 | 11.5±1.1 | 11±0.4 | **a vs. b**<br>**0.011**<br>a vs. c<br>0.0002 |
| **PTT (seconds)** | 31.2±4.6 | 30.9±4 | 29.5±3.3 | 0.595 |
| **D-Dimer (ng/mL)** | 3.9±6.9 | 1513±1474 | 3215±1247 | **a vs. b**<br>**0.028**<br>a vs. c<br><0.0001 |
| **Fibrinogen**<br>**(mg/dL)** | 662.7±167.5 | 693.9±186.3 | 566.2±103.4 | 0.195 |
| **Oxygen saturation**<br>**(%)** | 88.4±13.3 | 91.1±5.6 | 98.5±0.5 | **0.049** |
| **Diabetes** |  |  |  |  |
| Gestational | 2 | 3 | 1 | 0.614 |
| Type II |  |  |  |  |
| **Hypertension** | 1 | 2 | 0 | 0.375 |
| **Obesity** | 5 | 6 | 0 | **0.037** |
| **Gestational age (Weeks)** | -- | 31.4±5.7 | 35.3±4.8 | 0.069 |
| **1<sup>st</sup> trimester** | -- | 0 | 0 | ns |
| **2<sup>nd</sup> trimester** | -- | 2 | 1 | ns |
| **3<sup>rd</sup> trimester** | -- | 13 | 12 | ns |

Non-Pregnant COVID-19+ (NP-COVID-19+), Pregnant-COVID-19+ (P-COVID-19+), Pregnant-COVID-19- (P-COVID-19-). BMI; Body Mass Index. NLR; Neutrophil/Lymphocyte Ratio. Kruskal-Wallis test and Dunn's multiple comparisons test. Significant $p < 0.05$ . Media±SD.

**Table 2.** Frequency of COVID-19 signs and symptoms.

|  | **NP-COVID-19 +**<br>**(n=15)a** | **P-COVID-19 +**<br>**(n=13)b** | **P-COVID-19 -**<br>**(n=12)c** | ***p*** |
| --- | --- | --- | --- | --- |
| **Cough** | 10 | 9 | 0 | **0.001** |
| **Rhinorrhea** | 4 | 6 | 0 | **0.038** |
| **Odynophagia** | 5 | 3 | 0 | 0.092 |
| **Myalgia** | 10 | 8 | 0 | **0.001** |
| **Arthralgia** | 7 | 7 | 0 | **0.012** |
| **Anosmia** | 2 | 1 | 0 | 0.417 |
| **Dyspnea** | 11 | 6 | 0 | **0.0006** |
| **Diarrhea** | 3 | 1 | 0 | 0.202 |

Non-Pregnant COVID-19+ (NP-COVID-19+), Pregnant-COVID-19+ (P-COVID-19+), Pregnant-COVID-19- (P-COVID-19-). Fisher's exact test. Significant $p < 0.05$ .

### Blood sample collection

Our study is in accordance with the World Medical Association’s Declaration of Helsinki. After a patient agrees to participate in the study, healthcare personnel collected blood specimens in silicone coated, EDTA and heparinized tubes (BD Vacutainer, N.J, USA). Samples were processed immediately after collection. Whole blood count and serum was obtained to compare among groups.

### Leukocyte surface immunophenotype

Whole blood samples (50 µL) were incubated with titrated volumes of antibodies according to the following panel: all antibodies were from BioLegend, San Diego, CA, USA. Anti-CD45-PerCP (Clone:HI30), anti-CD3-AF647 (Clone:UCHT1), anti-CD14-PECy7 (Clone:M5E2), anti-CD16-FITC (Clone:3G8), anti-CD19-APC/Cy7 (Clone:HIB19), anti-CD73-PE (Clone:AD2), anti-CD39-BV421 (Clone:A1), anti-CD4-APC/Cy7 (Clone:OKT4), anti-CD8-PE/Dazzle594 or BV510 (Clone:SK1), anti-CD69-BV421 (Cone:FN50), anti-HLA-DR-AF488 o PE/Dazzle594 (Clone:L243), and for exclusion of dead cells Zombie Aqua fixable viability kit (BioLegend, San Diego, CA, USA) were used. After 15 minutes of incubation, erythrolysis was performed using FACS™ Lysing Solution (Cat. 349202, BD, San Jose, CA, USA). Samples were washed once with PBS 1x (1,500 rpm, 5 minutes, 4°C), and resuspended in PBS (100 µL). At least 30,000 leukocytes were acquired in a FACS Aria IIu flow cytometer (BD Biosciences, San José, CA, USA). The FACS files were analyzed with Infinicyt™ software 1.8 (Cytognos, Salamanca, Spain). Single cells were defined with FSC-A *vs*. FSC-H plot, and leukocytes were identified using a SSC *vs*. CD45 plot. Lymphocytes were gated as SSC^low^FSC^low^CD45^++^CD14^-^, monocytes as SSC^mid^FSC^mid^CD45^+^CD14^+^, and granulocytes-neutrophils as SSC^mid^FSC^mid^CD45^+^CD16^+^. Percentage and mean fluorescence intensity (MFI) of HLA-DR, CD69, CD39, CD73, CD32 and CCR5 positive cells were calculated.

### Cell culture

Whole blood (1 mL per well) was incubated (4 hours at 37°C with 5% CO2) alone in 24-well culture plates (Cat 13485, Costar, NY, USA), in the presence of human recombinant IL-6 (human rIL-6, 100 ng/mL), *Escherichia coli* O55: B5 Lipopolysaccharide (LPS 250 ng/mL, Cat. L2880, Sigma Aldrich, St. Louis, MO, USA), or Phorbol Myristate Acetate/Ionomycin (PMA 50 ng/mL, Ion 1mg/mL). In addition, two sets of samples were incubated, in the presence or absence of Brefeldin-A (Cat. 420601, BioLegend, San Diego, CA, USA). Afterwards, either intracellular phenotyping was performed, or supernatant was recovered and stored at -20°C until cytokine and chemokine assessment.

### Intracellular cytokine immunophenotype

After cell culture, whole blood samples (50 µL) were incubated with the following panel: antibodies from BioLegend, San Diego, CA, USA: anti-CD45-PerCP (Clone: HI30), anti-CD3-AF647 (Clone: UCHT1), anti-CD14-PECy7 (Clone: M5E2), anti-CD4-APC/Cy7 (Clone: OKT4), anti-CD8-PE/Dazzle594 or BV510 (Clone: SK1). After 15 min in the dark, blood was washed once with PBS (1mL) by centrifugation at 900×g for 5 min at RT, then Fixation buffer was added (100 μL, Cat: 420801, BioLegend, San Diego, CA, USA), and incubated for 20 min. Samples were then washed twice with 1 mL of Intracellular Staining Perm Wash buffer (Cat: 421002, BioLegend, San Diego, CA, USA); after the second wash, they were mixed with monoclonal antibodies against cytokines from BioLegend, San Diego, CA, USA: anti-TNFα-BV421 (Clone: MAb11), anti-IL-6-PE (MQ2-13A5), anti-IL-1β-FITC (Clone:JK1B-1), anti-IFNγ-BV421 (Clone:4S.B3), anti-IL-8a-PE (Clone: E8N1). For the exclusion of dead cells Zombie Aqua fixable viability kit (BioLegend, San Diego, CA, USA) was adjoined and incubated for 30 min dRT. Lastly, the mixture was washed once with PBS. At least 30,000 events were acquired in a FACS Aria IIu (BD, San Jose CA) flow cytometer. Analysis was performed using the Infinicyt™ Software 1.8.

### Serum or supernatant cytokine/chemokines concentration

Serum or cell culture supernatant was analyzed as follows; cytokines (IL-2, IL-4, IL-6, IL-10, TNF-α, IFN-γ, and IL-17a) and chemokines (CXCL8/IL-8, CXCL10/IP-10, CCL11/Eotaxin, CCL17/TARC, CCL2/MCP-1, CCL5/RANTES, CCL3/MIP-1a, CXCL9/MIG, CXCL5/ENA-78, CCL20/ MIP-3a, CXCL1/GROa, CXCL11/I-TAC and CCL4/MIP-1b) were determined using bead based immunoassays (CBA kit, Cat. 560484, BD PharMingen, San Diego, CA, USA; and LEGENDplex, Cat. 740003, BioLegend, San Diego, CA, USA). Log-transformed data were used to construct standard curves fitted to 10 discrete points using a 4-parameter logistic model. The concentration of each cytokine/chemokine was calculated using interpolations of their corresponding standard curves.

### Statistical analysis

Statistical analysis was performed using GraphPad Prism® version 7 software (GraphPad Software, San Diego, CA, USA). Non-parametric ANOVA test (Kruskall-Wallis test) with Dunn post-test were applied. Categorical variables were expressed as percentage number (%) and compared by Fisher’s exact test. A *p*<0.05 was considered as statistically significant.

## Results

To assess the immune profile in pregnant women infected with SARS-CoV-2, we analyzed; a) NP-COVID-19+, b) P-COVID-19+ and c) P-COVID-19-. Table 1 shows the clinical characteristics and laboratory values. No statistically significant difference was observed for age, BMI, respiratory rate, body temperature, Mean Arterial Pressure (MAP), hemoglobin, total leukocyte count, Neutrophil/Lymphocyte Ratio (NLR), total platelet count, serum glucose, serum creatinine, Partial Thromboplastin Time and fibrinogen between the NP-COVID-19+ or P-COVID-19-groups *vs*. P-COVID-19+. We did not observe any difference in the leukocyte count among groups, this is consistent after phenotyping analysis by flow cytometry (Table S1). Furthermore, the frequency of comorbidity (diabetes mellitus and systemic arterial hypertension) was similar among the groups. On the other hand, some clinical characteristics display differences, for example, heart rate was higher in P-COVID-19+ than in P-COVID-19-patients (p=0.048). Likewise, serum Lactate Dehydrogenase (LDH) concentration was higher in P-COVID-19+ than in P-COVID-19-group. Nevertheless, patients in the P-COVID-19-group maintain higher oxygen saturation levels than in COVID-19+ patients with or without pregnancy. Regarding serum D-dimer concentrations, higher magnitudes were reported in P-COVID-19+ than in NP-COVID-19+. It is worthy of highlighting that a SARS-CoV-2 infection does not increase the D-dimer concentration to the levels reported in physiological pregnancy. Similar gestational age, and number of cases per trimester were analyzed in pregnant women.

Table 2 shows the frequency of symptoms exhibited by patients with and without COVID-19. Despite gestation, patients with COVID-19 had a similar frequency of symptoms such as cough, myalgia, arthritis and anosmia. The most frequent symptoms in pregnant patients with COVID-19 were: cough, myalgia, and arthralgia.

Cellular immune response is essential against SARS-CoV-2 infection. Using flow cytometry, the proportion of leukocytes with an activated phenotype (HLA-DR+ or CD69+) or inflammatory regulators (CD39+ or CD73+) was determined in peripheral blood samples from the three groups by phenotyping, as described in the methods, and were compared among each other. Figure 1 shows the lower proportion of monocytes HLA-DR+ found in P-COVID-19+ compared to P-COVID-19-patients (p=0.006). Along the same line, a lower percentage of monocytes HLA-DR+ was observed in NP-COVID-19+ than in P-COVID-19-group (p=0.0003), while NP-COVID-19+ and P-COVID-19+ show similar percentage of monocytes HLA-DR+. In contrast, the percentage of CD4 or CD8 T lymphocytes that express HLA-DR or the early activation marker (CD69) were similar among groups (Table S2). Regarding CD39, the percentage of monocytes CD39+ was lower in the NP-COVID-19+ than in the P-COVID-19+ group (p=0.004), a lower percentage was noticed in NP-COVID-19+ than in P-COVID-19-group; however, this difference was not statistically significant (p = 0.663). A tendency in P-COVID-19+ women to express higher percentage of monocytes CD39+ compared to non-infected gestating women (p=0.084) was observed. CD39 expression was furthermore determine in B and T lymphocytes, noticing a higher percentage of B CD39+ cells in P-COVID-19+ and P-COVID-19-compared to NP-COVID-19+ group, but these differences were not statistically significant (p=0.127, p=0.487, respectively). On the other hand, T CD39+ cells are higher in NP-COVID-19+ than in P-COVID-19+ or P-COVID-19-groups, although these differences were not significant (p=0.201, p=0.057 respectively). With respect to CD73, it was found that monocytes CD73+ reached a higher percentage in NP-COVID-19+ than in P-COVID-19+ group (p=0.029), while, the NP-COVID-19+ and P-COVID-19-show a very similar percentage of monocytes CD73+ (p> 0.999). Despite pregnancy or COVID-19 similar percentages of CD73+ in B cells were detected. In contrast, the percentage of T CD73+ cells were lower in NP-COVID-19+ than in P-COVID-19-(p=0.038), but similarly low between COVID-19+ women regardless of their gestational status (p=0.227).

**Figure 1.**
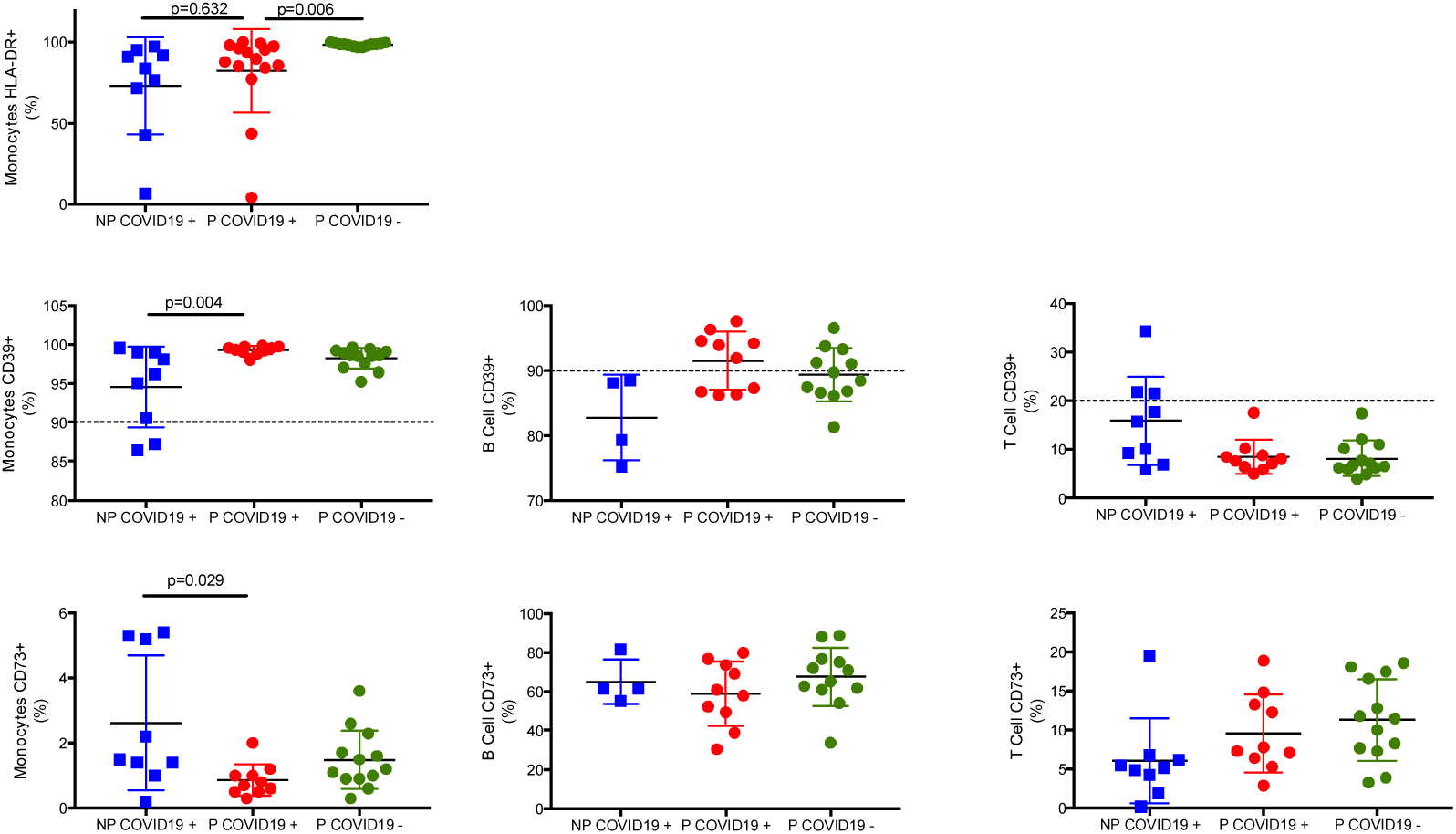
Surface marker on leukocytes. Whole blood cells were immunophenotype as described in methods. Results are expressed as mean±SD. Significance value was *p*<0.05. Kruskal-Wallis and Dunn’s multiple comparisons test was calculated. Non-Pregnant COVID-19 positive (NP-COVID-19+, n=4-9). Pregnant-COVID-19 positive (P-COVID-19+, n=10-15). Pregnant-COVID-19 negative (P-COVID-19-, n=12-13). Dot line indicates the percentage of monocytes, B and T cells that constitutively express CD39 (26).

The proportion of lymphocytes and monocytes that synthesize cytokines IL-6, IFN-γ or IL-1β was determined in P-COVID-19+ patients. Whole blood samples were cultivated 4 hours in absence or presence of polyclonal stimuli (LPS 250 ng/mL, PMA/Ion, 50ng/mL/1mg/mL) or human rIL-6 (100 ng/mL). Figure 2 shows the results of this functional assay. The percentage of CD4 or CD8 T lymphocytes expressing IL-6 or IFN-γ was less than 5% in the groups (Figure 2a, d, g, j), and human rIL-6 did not increase this percentage in CD4 or CD8 T lymphocytes (Figure 2b, e, h, k). The stimulus with PMA/Ion did increase the percentage of CD4 IL-6+ T lymphocytes in the P-COVID-19+ group with respects to the P-COVID-19-group. However, this effect was not seen in the CD8+ IL-6+ T lymphocytes (Figure 2c, i). In addition, PMA/Ion stimulus increased the percentage of CD4 and CD8 IFN-γ+ cells in total blood of pregnant women with and without COVID-19 (Figure 2f, l), but statistical significance is only reached in CD8 IFN-γ+ T lymphocytes between the P-COVID-19+ and P-COVID-19-groups (p=0.019, Figure 2l).

**Figure 2.**
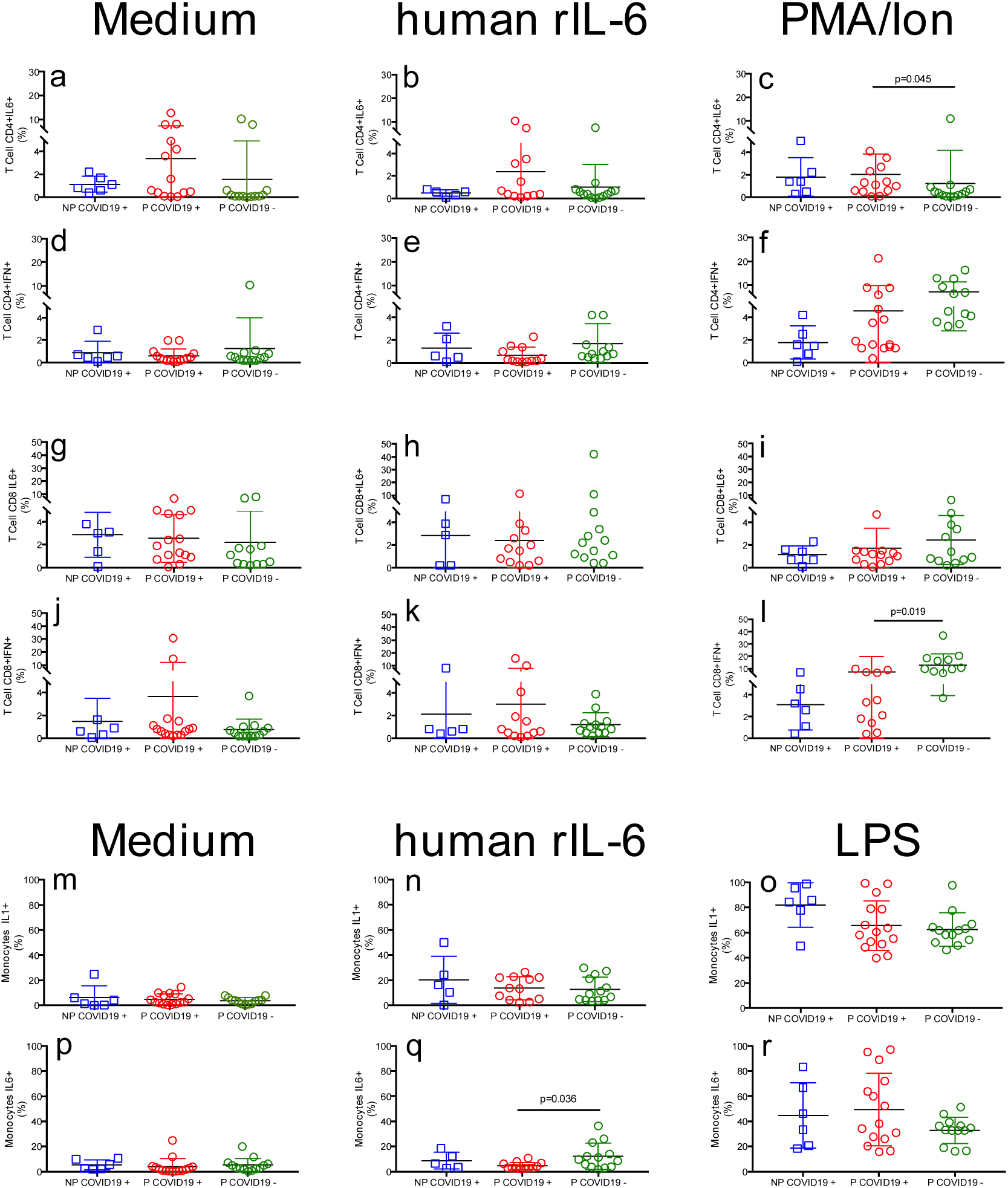
Percentage of cytokine-positive leukocytes. Whole blood cells were immunophenotype as described in methods. Results are expressed as mean±SD. Significance value was *p*<0.05. Kruskal-Wallis and Dunn’s multiple comparisons test was calculated. Non-Pregnant COVID-19 positive (NP-COVID-19+, n=5-6). Pregnant-COVID-19 positive (P-COVID-19+, n=12-15). Pregnant-COVID-19 negative (P-COVID-19-, n=12-13).

By the same token, the percentage of monocytes IL-1+ or IL-6+ is less than 10% in the groups (Figure 2m, p). The stimulus with human rIL-6 only increased the percentage of monocytes IL-6+ in the P-COVID-19-group with respect to P-COVID-19+ (Figure 2q), but the percentage of monocytes IL-1+ was similar among groups (Figure 2n). Incubation with LPS increased the proportion of monocytes IL-1+ and IL-6+ in the three groups (Figure 2o, r), with no significant differences among groups. A lower percentage of monocytes IL-1+ was observed in pregnant women with and without COVID-19 than in NP-COVID-19+(Figure 2o, p=0.314 and p=0.213 respectively). In addition, the percentage of monocytes IL-6+ in patients with COVID-19+ with and without pregnancy was similar (Figure 2r). After the LPS challenge, we documented a lower percentage of monocytes IL-6+ in P-COVID-19-than in P-COVID-19+ group, even so, this difference is not statistically significant (p=0.724).

To determine the serum concentration of cytokine/chemokine in pregnant women with COVID-19+, samples were collected and compared with controls (NP-COVID-19+ and P-COVID-19-). The MIP1b, TNF-α, IL-6 and IL-4 concentration is higher in P-COVID-19+ than in P-COVID-19- (Figure 3a, b, d, e). Also, the TNF and IL-4 concentration is higher in P-COVID-19+ than in NP-COVID-19+ (Figure 3d, e). Other cytokine/chemokine show higher concentration in P-COVID-19+ than in P-COVID-19-, such as CXCL10 (IP10) and IL-2, although these do not reach a statistically significant difference (Figure 3c and f, p = 0.456, and p=0.447, respectively). Figure S1 shows other cytokine/chemokine that show similar concentration between and among groups, these includes CXCL8, CCL11, CCL17, CCL2, CCL5, CCL3, CXCL9, CXCL5, CCL23, CXCL1, CXCL11, IL-17a, IFN-γ and IL-10.

**Figure 3.**
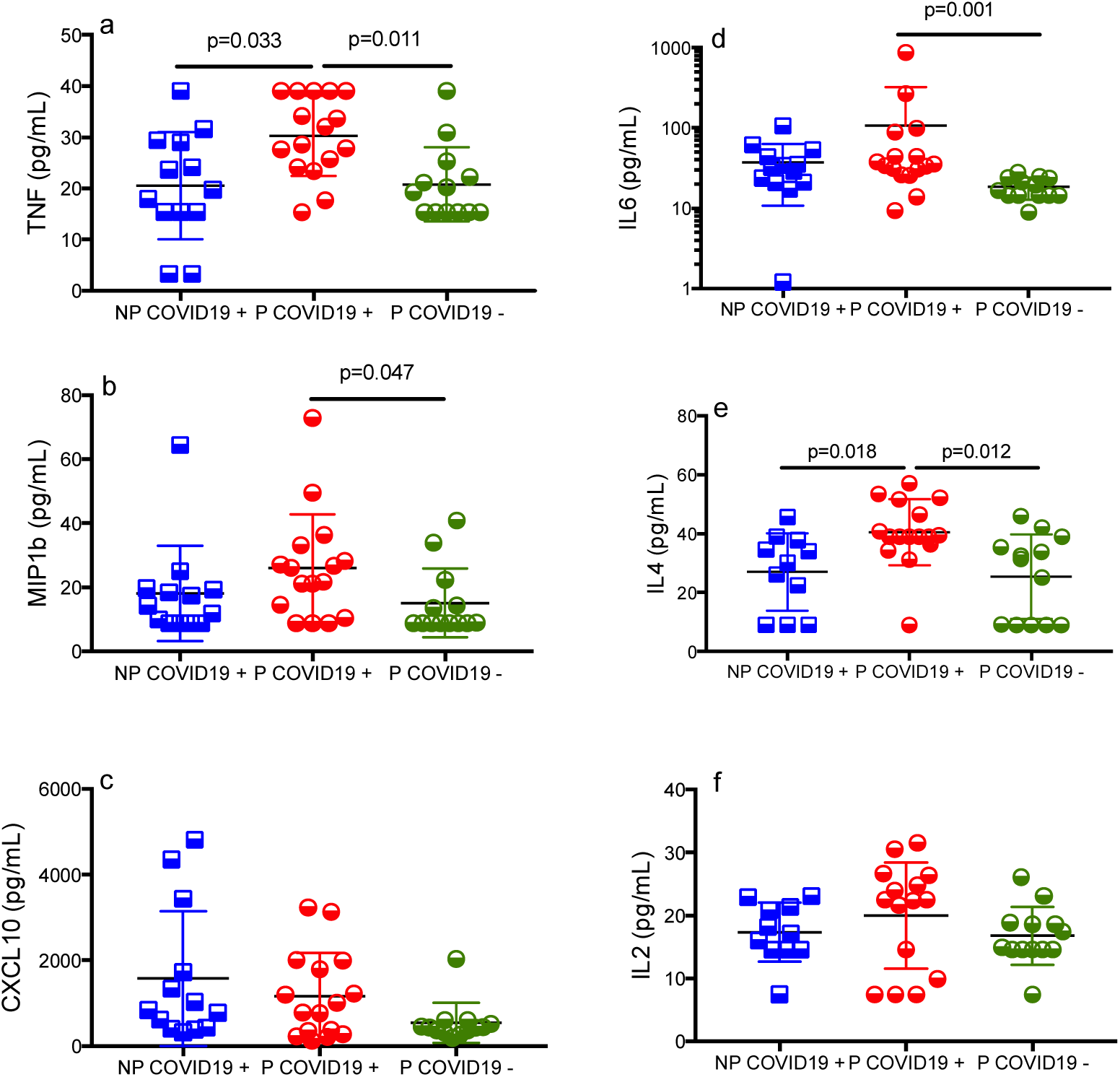
Cytokine/Chemokine serum concentration. Serum was isolated and cytokine/chemokines concentration was determined using bead-based immunoassays as described in methods. Results are expressed as mean±SD. Significance value was *p*<0.05. Kruskal-Wallis and Dunn’s multiple comparisons test was calculated. Non-Pregnant COVID-19 positive (NP-COVID-19+, n=13). Pregnant-COVID-19 positive (P-COVID-19+, n=14-15). Pregnant-COVID-19 negative (P-COVID-19-, n=13).

To know whether leukocytes from pregnant women with COVID-19 can be activated by polyclonal confrontation and lead to cytokine/chemokine response, whole blood was challenged; Figure 4 shows the concentration in supernatant. After 4 hours of culture without stimulation (whole blood only), the concentration of TNF-α, IFN-γ, CCL3, CCL4, IL-17a, CCL23, CXCL8 and IL-10 was similar among groups. Human rIL-6 induced a higher concentration of TNF-α, CCL3 and CCL4 (Figure 4e, f, d, h), but only TNF-α and CCL3 reached a difference statistically significant (Figure 4a, b). Also, LPS induced a higher concentration of TNF-α, CCL3, CCL4, CCL23 and CXCL8 in all groups, these differences reached high significance (p<0.0001) when contrasted with their respective pair without stimulus (in example; NP-COVID-19+ whole blood *vs*. P-COVID-19+ LPS). Finally, the stimulation with PMA/Ion induced a higher concentration of CCL3, CCL4, IL-17a, CCL23, CXCL-8 and IL-10 in all groups rather than in their respective pair without stimulus (in example; NP-COVID-19+ WB vs. P-COVID-19+ PMA/Ion). The PMA/Ion did not induce an increase IFN-γ response in all groups, especially in NP-COVID-19+ group, however, the IFN-γ concentration was higher in the pregnancy groups with and without COVID-19 compared to NP-COVID-19+ group, although no statistically significant difference was achieved (p> 0.9, p> 0.9 respectively). Figure S2 shows that the polyclonal stimulus did not induce an increase response of CXCL10, CCL11, CCL17, CCL2, CCL5, CXCL9, CXCL5, CXCL1, CXCL11, IL-10, IL-4 and IL-2.

**Fig 4.**
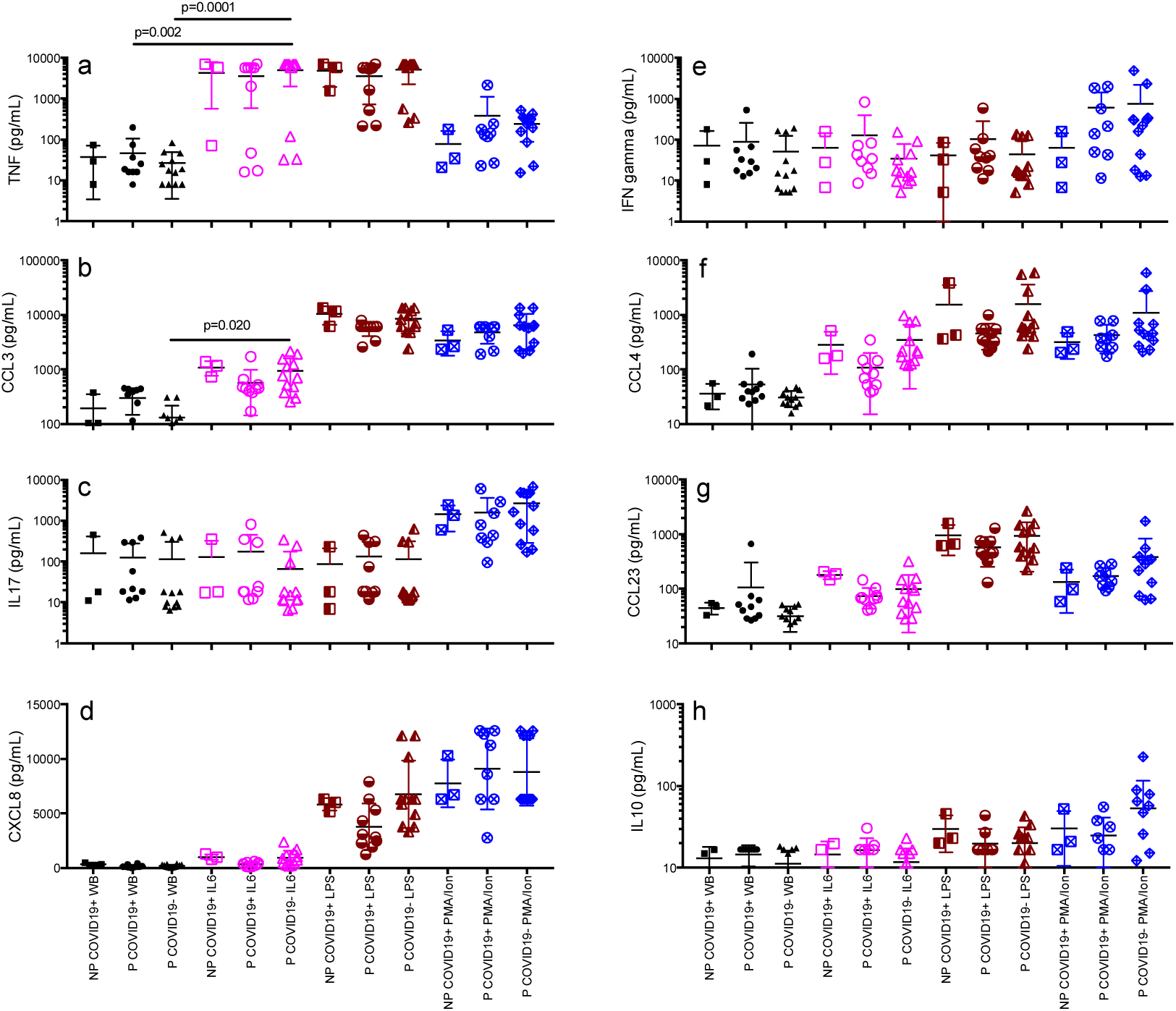
Cytokine/Chemokine response after 4 hours of culture with polyclonal stimulus in pregnant and non-pregnant women with or without COVID-19. Supernatant was collected and cytokines/ chemokines concentration was determined using bead-based immunoassays as described in methods. Results are expressed as mean±SD. Significance value was *p*<0.05. Kruskal-Wallis and Dunn’s multiple comparisons test was calculated. Non-Pregnant COVID-19 positive (NP-COVID-19+, n=3). Pregnant-COVID-19 positive (P-COVID-19+, n=8-10). Pregnant-COVID-19 negative (P-COVID-19-, n=12). WB, Whole Blood.

## Discussion

During pregnancy, the immune system is highly regulated. Multiple mechanisms of immune tolerance develop during gestation, and favor physiological progress in reproduction (6). Any viral infection poses a high risk of embryo/fetal-maternal morbi-mortality, as a result of deregulated cellular and humoral immune response immunity, especially during SARS-CoV-2 infection. Villar et al, reported a greater probability of morbidity and mortality in pregnant women with COVID-19 than in pregnant women without the disease (2). However, how the immune response is involved is unknown in great detail. Herein we explored certain cellular and humoral characteristics that may enhance the understanding of the immuno-pathophysiology in pregnant women with COVID-19.

Pregnant women with COVID-19 were analyzed and compared with non-pregnant women with COVID-19 or those during physiological pregnancy. NP-COVID-19+ and P-COVID-19+ groups were similar in several clinical characteristics such as, BMI, respiratory and heart rate, temperature, MAP, hemoglobin, total platelet count, serum glucose, serum creatinine, LDH and fibrinogen (Table 1). These results indicate that the series of cases analyzed had comparable clinical status. However, some differences between non-pregnant and pregnant women to highlight are: a higher serum urea concentration, the longer PT time and the lower concentration of D-Dimer in NP-COVID-19+ than in P-COVID-19+ or P-COVID-19-. Such differences could be due to the physiological change in the coagulation system during pregnancy (27), and not due to the SARS-CoV-2 infection, moreover, there is a very few reports of COVID-19 coagulopathy during pregnancy(28). A high D-Dimer concentration has been previously reported in the general population as well as in pregnant women with COVID-19 correlating its high levels with fatal outcomes (29-31). Interestingly, within this analysis there was no higher concentration of D-Dimer in P-COVID-19+ compared with P-COVID-19-patients suggesting that, unlike in the general population, D-Dimer concentrations in pregnant women are not necessarily indicative of severity or thromboembolic risk but rather a physiological state of gestation by itself (32). It is necessary to increase the number within a longitudinal study to assess the usefulness of D-Dimer concentration as a severity predictor factor in pregnant COVID-19+ women. Additionally, the frequency of symptoms was similar between NP-COVID-19 and P-COVID-19 (Table 2), indicating a homogeneous clinical presentation, allowing the identification of cellular and humoral characteristics in response against COVID-19 in the presence or absence of pregnancy.

Our study shows that the percentage of monocytes HLA-DR+ is lower in COVID-19+ women with and without pregnancy than in P-COVID-19- (Fig 1). Also, a lower percentage has been observed in septic patients with critical condition or fatal outcome (33, 34), suggesting that this characteristic could be a helpful biomarker in COVID-19. The lower percentage of monocytes HLA-DR+ in COVID-19 could be the way to downregulate the immune response by SARS-CoV-2 and to evade the immunity, or the way that the immune response control activation and avoid over-stimulation. Our results suggest that COVID-19 does not accentuate the low percentage of monocytes HLA-DR+, which would indicate that pregnancy does not limit the activation of monocytes in peripheral blood upon a SARS-CoV-2 infection. More analysis is required to know the biological significance of lower percentage of monocytes HLA-DR+ in COVID-19. Furthermore, a high percentage of lymphocytes CD69+ is reached in AH1N1 influenza, another viral challenge that leads to unregulated inflammation in pregnant women (35). Likewise, we found a higher percentage of CD69+ cells in both CD4 and CD8 cells in P-COVID-19+ than in NP-COVID-19+ and P-COVID-19-without reaching a statistical significance (Table S2). This indicates a higher level of activation despite the multiple mechanisms to ensure an immunotolerance during pregnancy.

The expression of CD39 and CD73 on leukocytes may be an important mechanism for resolving SARS-CoV-2 infection and COVID-19 disease. CD39 and CD73 are ectoenzymes that sequentially metabolize ATP to adenosine (26, 36), thus controlling inflammation through this alarmine, leading to an adenosine induced anti-inflammatory response (26). Pregnant women with or without COVID-19 maintain higher percentages of monocytes CD39+, and lower percentage of CD73+ than in NP-COVID-19+ (Fig 1), suggesting that pregnant women control inflammation through monocytes CD39+/CD73+. These could be a potential marker to monitor the evolution of COVID-19. The function of CD39/CD73 is not limited to monocytes, however, our results indicate that the percentage of B or T cells CD39+ or CD73+ is not significantly modified by the effects of pregnancy or COVID-19 infection (Figure 1).

Activated leukocytes are a potential source of pro-inflammatory or regulatory cytokines in peripheral blood of COVID-19 patients. We determined the percentage of leukocytes IL-6+, IFN-γ+ or IL-1β+ after 4 hours of culture with or without polyclonal stimulation. Lymphocytes T CD4+ IL-6+ or IFN-γ+, CD8+IL-6+ or IFN-γ+ and monocytes IL-1β+ or IL-6+ did not reach more than 5% of circulating cells, indicating a low baseline of circulating cytokine producing leukocytes. After being stimulated with human rIL6, there was no significant increase of IL-1β, IL-6 or IFN-γ producers in lymphocytes and monocytes, indicating that the IL-6 in serum of COVID-19 patients could have a limited stimulus to increase the synthesis of pro-inflammatory cytokines from circulating leukocytes. Whole blood stimulation LPS or PMA/Ion increases the percentage of lymphocytes T CD4+ and CD8+ and monocytes IFN-γ+ or IL-1β+ and IL-6+, indicating that these leukocytes are not anergic and retain the ability to synthesize cytokines both in the presence and absence of pregnancy and COVID-19. However, there is a trend to increase the percentage of monocytes IL-6+ and decrease the percentage of lymphocytes CD4+IFN-γ+ and CD8+IFN-γ+ in patients with COVID-19. This suggests that both women with or without pregnancy develop quite similar defenses to face COVID-19, increasing IL-6 and limiting IFN-γ response.

The cytokine storm induced by COVID-19 could be greater in pregnant women, our study shows that some cytokines (TNF-α, IL-6 and CCL3) reach a higher concentration in serum of NP-COVID-19+ than in P-COVID-19+ patients, although it was only significantly higher for TNF-α (Figure 3a). Interestingly, we also found a highest concentration of IL-4 in the P-COVID-19+ group with a statistically difference in NP-COVID-19+ (p=0.01) and P-COVID-19- (p=0.01). Despite pregnancy, results indicate a similar pro-inflammatory profile in COVID-19+ patients, which could be regulated by IL-4 in pregnant women. Some reports show that the concentration of CXCL10 chemokine is associated with a poor prognosis in COVID-19+ patients (20). In contrast, this study found a lower concentration in P-COVID-19+ patients, which would favor the immune response in pregnancy.

After analysis of the basal and leukocyte response to polyclonal stimulation, the basal concentration of cytokines is similar among groups (Figure 4), indicating that, despite COVID-19, peripheral leukocytes have a similar potential to produce cytokines. This suggests that production of cytokines in COVID-19 could depend of an alternate source such as endothelial cells. Human rIL-6 stimulus caused an increase in some cytokines such as TNF-α, CCL3 and CCL4, indicating that IL-6 favors synthesis of some cytokines but not an entire cytokine storm. In addition, the response in COVID-19 to IL-6 seems to be similar in the presence or absence of pregnancy, indicating that pregnancy not necessarily aggravates the pro-inflammatory responses in COVID-19. To explore if leukocytes response is limited by COVID-19, we performed a LPS or PMA/Ion stimulus, resulting in a clear pro-inflammatory response with cytokines such as, TNF-α, CCL3, CCL4, IL-17a, CCL23 and CXCL8 in supernatant, indicating that leukocytes are not anergic. It has been proposed that an immunosuppression rather than a hyper-cytokine response in COVID-19 could support the pathophysiology (37). However, these results indicate that peripheral blood leukocytes from pregnant women with COVID are capable of expressing a similar response as a healthy pregnant woman. The main limitation of the present study is the compact number of patients. A greater number of observations are required to reach a final and more representative conclusion. However, the reproducibility and consistency of these results back our analysis. Pregnant women with or without COVID-19 control inflammation through monocytes CD39+/CD73+ maintaining higher percentages of monocytes CD39+ and lower percentage of CD73+ than NP-COVID-19+ patients. Hence, CD39/CD73 is a potential marker to monitor the evolution of COVID-19. On the other hand, unlike in the general population, D-Dimer concentrations in pregnant women are not necessarily indicative of severity or thromboembolic risk but rather a physiological state of gestation by itself. These findings help the focus for future studies on the immune profile in pregnant women with COVID-19, provides evidence about the functional immunological profile in pregnant women with COVID-19 and enriches the knowledge on the immune response that occurs during pregnancy.

## Contributors

ACV and CLM conceived and designed the study, and contributed to data analysis. ACV wrote the first version of manuscript.MGE, JCBG, LAAP, EFO, OMA, GMLGA, contributed with a critical revision of the report. MGE, JCBG, BZB, RCdLB, AHVC, AEP, RRMdO, FCS, LARG, GFP, OMA, GMLGA contributed with the clinical evaluation of patients and supervision of medical treatments and patient care. ACV, LAAP, EFO, GLCR, PMC, MTGR, JLPC, VRA, RLMS, ACC, ESR, MESR and DSM, contributed with data acquisition, analysis and/or interpretation. All authors reviewed the final version. CLM reviewed and approved the final version.

## Acknowledgements

This project was supported by the Mexican National Research Council (CONACyT), (Project No. 313494 awarded to CLM). The authors extend a gratitude to the staff at the Specialties Hospital, National Medical Center “XXI Century”, Gynecology & Obstetrics Department in the General Hospital of Mexico “Dr. Eduardo Liceaga” and Gynecology & Obstetric Hospital No. 4 UMAE “Dr. Luis Castelazo Ayala”.

---

**Table S1.** Percentage of leukocytes after 4 hours of medium culture in pregnant and non-pregnant women with or without COVID-19.

| Leukocyte (%) | NP-COVID-19+<br>(n=6-9)a | P-COVID-19+<br>(n=15)b | P-COVID-19-<br>(n=13)c | *p* |
| --- | --- | --- | --- | --- |
| **Lymphocyte** | 15.8±11.4 | 19.4±9.5 | 28.2±7.8 | **0.031** |
| T cell CD4+ | 55.3±8.4 | 54.5±8.9 | 59.7±7.4 | 0.206 |
| T cell CD8+ | 35.5±9.3 | 37.7±9.9 | 34.1±6.0 | 0.533 |
| T cell CD4+CD8+ | 0.6±0.4 | 1.2±1.9 | 0.9±1.2 | 0.801 |
| **Monocytes** | 2.4±1.3 | 2.5±1.7 | 3.4±1.8 | 0.359 |
| **Granulocytes** | 81.5±12.5 | 77.8±10.4 | 68.1±9.4 | 0.058 |

Non-Pregnant COVID-19+ (NP-COVID-19+), Pregnant-COVID-19+ (P-COVID-19+), Pregnant-COVID-19- (P-COVID-19-). Fisher's exact test. Significant $p<0.05$ . Viability >90%.

**Table S2.** Surface of activation markers on leukocytes in pregnant and non-pregnant women with or without COVID-19.

| Leukocyte (%) | NP-COVID-19+<br>(n=4-9)a | P-COVID-<br>19+<br>(n=10-15)b | P-COVID-19-<br>(n=12-13)c | *p* |
| --- | --- | --- | --- | --- |
| **T cell CD4+CD69+** | 0.9±0.4 | 4.9±13.8 | 0.4±0.2 | a vs. c<br>0.020 |
| **T cell CD8+CD69+** | 5.1±4.2 | 5.3±4.6 | 2.4±1.3 | 0.076 |
| **T cell CD4+HLA-DR+** | 5.2±2.5 | 5.4±3.6 | 6.2±2.0 | 0.509 |
| **T cell CD8+HLA-DR+** | 10.5±9.5 | 11.2±7.2 | 14.8±6.1 | 0.121 |

Non-Pregnant COVID-19+ (NP-COVID-19+), Pregnant-COVID-19+ (P-COVID-19+), Pregnant-COVID-19- (P-COVID-19-). BMI; Body Mass Index. NLR; Neutrophil/Lymphocyte Ratio. Kruskal-Wallis test. Significant $p<0.05$ . Media±SD.

**Table S3.** Cytokine-positive leukocyte percentage after 4 hours of polyclonal stimulus in pregnant and non-pregnant women with or without COVID-19.

| Leukocyte (%) | NP-COVID-19<br>+<br>(n=5-6)a | P-COVID-19 +<br>(n=12-15)b | P-COVID-19<br>-<br>(n=13)c | *p* |
| --- | --- | --- | --- | --- |
| **T cell CD4+TNF+** |  |  |  |  |
| **Medium** | 0.4±0.2 | 0.9±1.2 | 0.7±1.4 | 0.222 |
| **IL-6** | 3.5±6 | 0.4±0.4 | 0.6±0.6 | 0.290 |
| **PMA/Ion** | 8.4±7.3 | 9.9±8.2 | 14.6±16.7 | 0.692 |
| **T cell CD8+TNF+** |  |  |  |  |
| **Medium** | 1.4±1.0 | 3.7±8.1 | 0.5±0.7 | 0.118 |
| **IL-6** | 2.3±4.3 | 1.3±3.0 | 0.6±1.0 | 0.337 |
| **PMA/Ion** | 4.6±2.5 | 8.3±8.1 | 11.2±9.6 | 0.210 |
| **Monocyte TNF+** |  |  |  |  |
| **Medium** | 5.4±5.2 | 3.7±4.1 | 1.9±2.1 | 0.218 |
| **IL-6** | 4.6±2.7 | 4.5±4.8 | 4.4±3.2 | 0.814 |
| **LPS** | 38.6±30.1 | 43.0±28.9 | 21.2±12.9 | 0.105 |
| **Monocyte IL-8+** |  |  |  |  |
| **Medium** | 14.1±10.6 | 13.9±9.6 | 22.3±12.5 | 0.165 |
| **IL-6** | 24.3±15.6 | 17.6±12.6 | 26.1±14.7 | 0.274 |
| **LPS** | 44.4±35.7 | 50.4±26.7 | 39.6±16.2 | 0.540 |

NP-COVID-19+; Non-Pregnant COVID-19+. P-COVID-19+; Pregnant-COVID-19+. P-COVID-19-; Pregnant-COVID-19-. Fisher's exact test. Significant $p<0.05$ .

**Figure S1.**
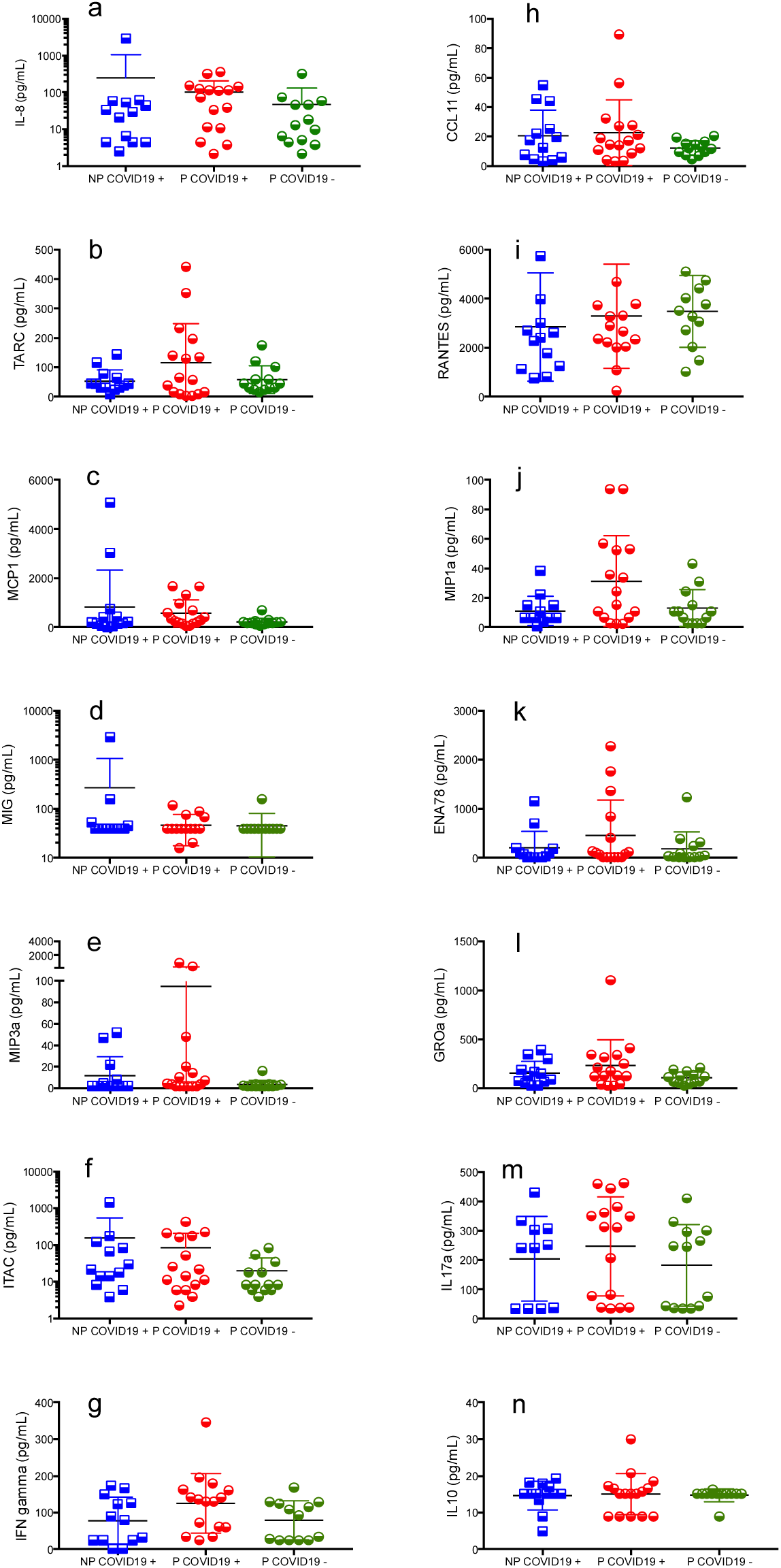
Similar Cytokine/Chemokine serum concentration in pregnant and non-pregnant women with or without COVID-19. Serum was isolated and cytokine/chemokines concentration was determined using bead-based immunoassays as described in methods. Results are expressed as mean±SD. Significance value was *p*<0.05. Kruskal-Wallis and Dunn’s multiple comparisons test was calculated. Non-Pregnant COVID-19 positive (NP-COVID-19+, n=13). Pregnant-COVID-19 positive (P-COVID-19+, n=15-16). Pregnant-COVID-19 negative (P-COVID-19-, n=13).

**Figure S2.**
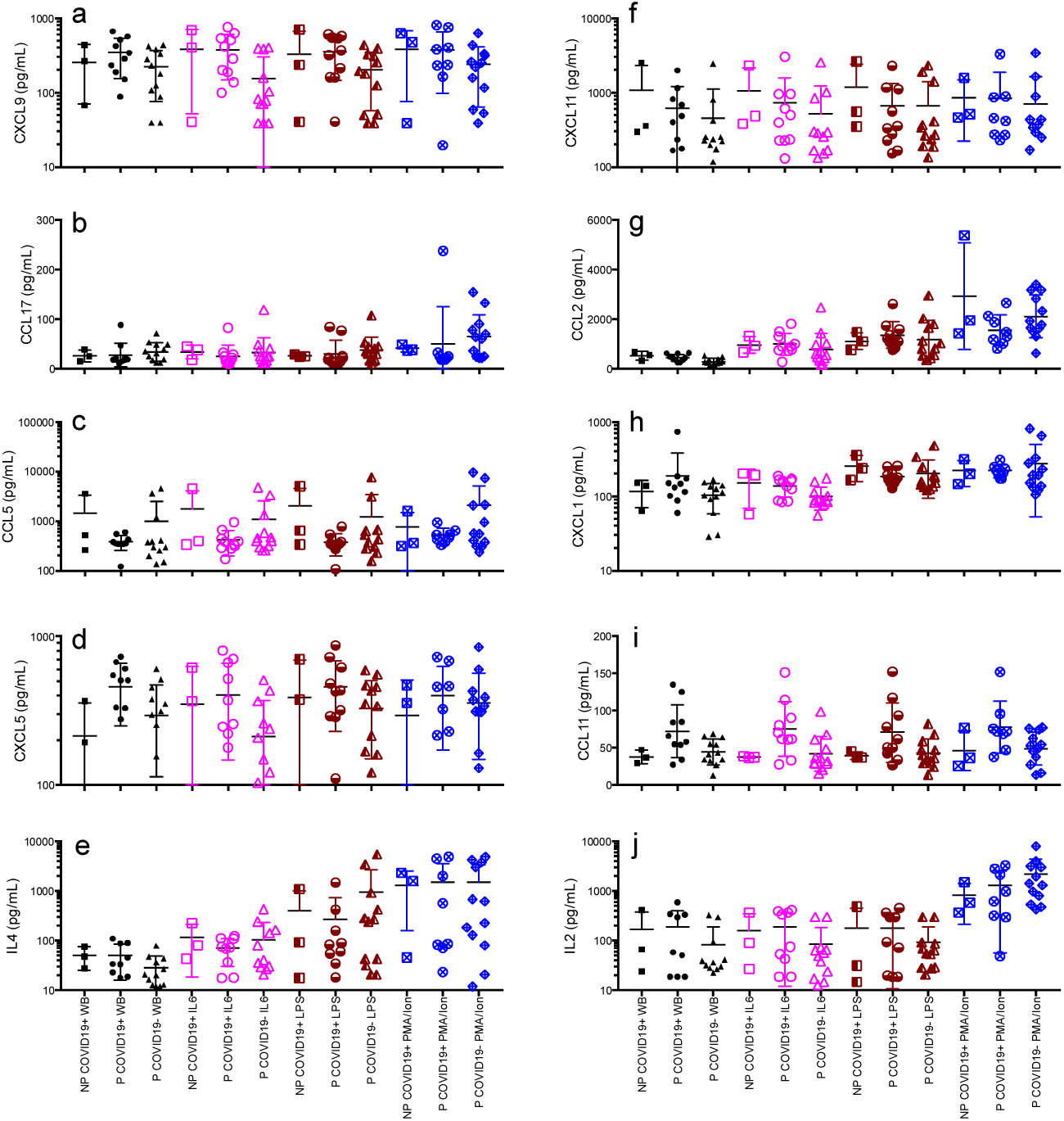
Similar Cytokine/Chemokine response after 4 hours of culture with polyclonal stimulus in pregnant and non-pregnant women with or without COVID-19. Supernatant was collected and cytokines/ chemokines concentration was determined using bead-based immunoassays as described in methods. Results are expressed as mean±SD. Significance value was *p*<0.05. Kruskal-Wallis and Dunn’s multiple comparisons test was calculated. Non-Pregnant COVID-19 positive (NP-COVID-19+, n=3). Pregnant-COVID-19 positive (P-COVID-19+, n=9-10). Pregnant-COVID-19 negative (P-COVID-19-, n=12). WB, Whole Blood.

